# che*C*k*OVER*: An open framework and AI-ready global crayfish database for next-generation biodiversity knowledge

**DOI:** 10.64898/2025.12.29.696807

**Authors:** Lucian Pârvulescu, David Livadariu, Victor I. Bâcu, Constantin I. Nandra, Teodor T. Stefănut, World of Crayfish® Contributors

## Abstract

**Background:** Species occurrence records represent the backbone of biodiversity science, yet their utility is often limited to spatial analyses, distribution maps, or presence-absence models. Current biodiversity infrastructures rarely provide computational formats directly usable by modern artificial intelligence (AI) systems, such as large language models (LLMs), which increasingly mediate scientific communication and knowledge synthesis. Open frameworks that convert biodiversity occurrences into structured, machine-accessible, provenance-rich knowledge are therefore essential—particularly those enabling rapid integration of new records, near real-time generation of spatial metrics, and production of both human-interpretable reports and AI-consumable outputs. Such capabilities substantially reduce latency between data acquisition and decision support, while ensuring biodiversity knowledge remains traceable and verifiable in AI-mediated workflows.

**Results:** We introduce cheCkOVER, an open framework that converts raw species occurrence datasets into standardized, API-ready, multi-layered outputs: biogeographic descriptors, dynamic distribution maps, summary metrics, and structured JSON geo-narratives following a canonical template. The framework stratifies processing by population origin (indigenous vs. non-indigenous), enabling IUCN-aligned conservation metrics while simultaneously tracking invasion dynamics. Each output embeds standardized citation metadata ensuring full provenance traceability. We applied the pipeline to 111,729 validated crayfish (Astacidea) occurrence records from 465 species, generating comprehensive species packages including indigenous-range classifications (171 endemic, 287 regional, 5 cosmopolitan taxa) and non-indigenous range tracking for 30 invasive species. This proof-of-concept demonstrates how the framework transforms minimal datapoints—validated species occurrences—into interoperable knowledge consumable by both humans and computational systems. The JSON outputs are optimized for retrieval-augmented generation, enabling AI systems to dynamically access and cite biodiversity knowledge with explicit source attribution.

**Conclusions:** che*C*k*OVER* is taxon-agnostic and establishes a reproducible pathway from biodiversity occurrences to narrative-ready, AI-interoperable knowledge with immediate public utility via the World of Crayfish® platform (https://world.crayfish.ro/), where each species page integrates structured outputs. The open-source framework (GPL-3) combines a generalizable processing pipeline with taxon-specific knowledge products, enabling flexible reuse across conservation research, policy reporting, and AI-driven applications. This minimalist-to-complex design extends the reach of biodiversity data beyond traditional analyses, positioning occurrence repositories as active knowledge engines for next-generation biodiversity informatics.

**Significance statement:** Biodiversity infrastructures remain underused by modern AI systems despite their central role in science and society. che*C*k*OVER* embodies a minimalist-to-complex paradigm: from the validated geographic occurrence of a species—a datapoint often perceived as trivial—it derives structured, multi-layered outputs linking distribution, conservation status, and standardized geographic indicators. These outputs are natively optimized for retrieval-augmented generation and other machine-consumable workflows, enabling AI systems to dynamically access and cite biodiversity knowledge with maintained provenance beyond their pre-training corpora. Using a global crayfish dataset as proof-of-concept, we demonstrate how raw occurrence records can scale into rich, interoperable biogeographic knowledge products with immediate value for both human experts and computational systems. This positions biodiversity databases as critical knowledge engines for next-generation science, policy, and societal decision-making, providing standardized outputs directly incorporable into conservation evaluation workflows where transparent, reproducible, and provenance-rich occurrence-based metrics are essential.

## Introduction

Human societies have always relied on knowledge to guide decisions and shape progress [1–3]. From antiquity to the digital era, science has served as a trusted source of verified information [4], while new technologies have multiplied the ways knowledge flows into public and policy domains [5,6]. Today, artificial intelligence (AI), particularly large language models (LLMs), is emerging as a dominant mediator of communication, offering rapid access to information across disciplines and audiences [7–9].

Species distribution records are highly valuable for numerous purposes, from scientific research [10,11] and understanding species communities [12,13], to conservation planning and management [14], and ultimately to certain economic considerations [15]. In particular, the quality of stored and delivered biodiversity knowledge is essential [16]. Nature underpins human well-being [17,18], and its understanding depends on reliable data about both species and the habitats they occupy [19,20]. Biodiversity knowledge must not only be abundant, but also accurate, transparent and easy to integrate into decision-making [21]. In this domain, misinformation or poorly contextualized data can directly affect conservation outcomes, policy priorities, and public trust [22,23].

However the speed and scale of LLMs analyses come at a cost: LLMs generate fluent but often vague, erroneous, or unverifiable outputs [24,25]. Their training depends on heterogeneous and inconsistently curated data, leaving a fundamental gap between scientific knowledge production and the AI systems that now mediate it [26]. Simultaneously, scientific data are expanding at unprecedented rates [27]. Advances in modeling, genomics, and ecological monitoring—together with citizen science—have produced millions of biodiversity records and traits [28–30]. Dedicated infrastructures and portals have proliferated, enabling access to vast repositories of occurrence data, images, and observations [31]. Yet the value of these resources remains underexploited [32,33]. These resources typically generate outputs that AI systems cannot directly consume, leaving only fragments of biodiversity knowledge contextualized in their reporting [34,35].

While humans digest information through narratives and visualizations [36,37], AI requires structured inputs: machine-readable formats (e.g., JavaScript Object Notation – JSON) annotated with explicit metadata and provenance [38–40]. To enable the full integration of biodiversity science in the AI era [41,42], infrastructures must evolve from human-facing repositories into dual-purpose systems [43–45]. Here we present che*C*k*OVER*, the first generalizable framework designed to fill this gap. At its core, che*C*k*OVER* deliberately returns to the roots of faunistics—validated raw occurrence records without prior interpretation—and transforms them into standardized, multi-layered outputs that are both human-readable and LLM-digestible. Using global crayfish (Astacidea) occurrence data [10] as a proof of concept, we illustrate how the framework scales minimal datapoints into interoperable, machine-readable knowledge packages that distinguish indigenous from non-indigenous populations while maintaining unified outputs for species assessment. These outputs are structured to support retrieval-augmented and other machine-consumable workflows, positioning biodiversity infrastructures to interface effectively with emerging AI technologies. In this way, che*C*k*OVER* reframes occurrence repositories as future-ready knowledge engines, enabling broader reuse, clearer attribution, and improved integration across scientific, conservation, and societal domains.

## Methods

### Framework description

che*C*k*OVER* is built as a modular architecture that transforms validated biodiversity occurrence records into standardized, multi-layered knowledge packages. Each package is generated at the species level and integrates visual, quantitative, and narrative components, ensuring outputs are both human-readable and machine-interoperable. The name cheCkOVER reflects the framework’s systematic checking of occurrences over various boundaries, signaling a step toward consolidated biodiversity knowledge infrastructure.

The workflow is organized around five conceptual components (Figure 1), internally implemented through seven sequential processing phases that ensure methodological rigor while accommodating complex distributional patterns. This hybrid design balances conceptual clarity with implementation flexibility, enabling the framework to handle species occurring exclusively as native populations, exclusively as introduced populations, or in both contexts simultaneously.

**Figure 1.**
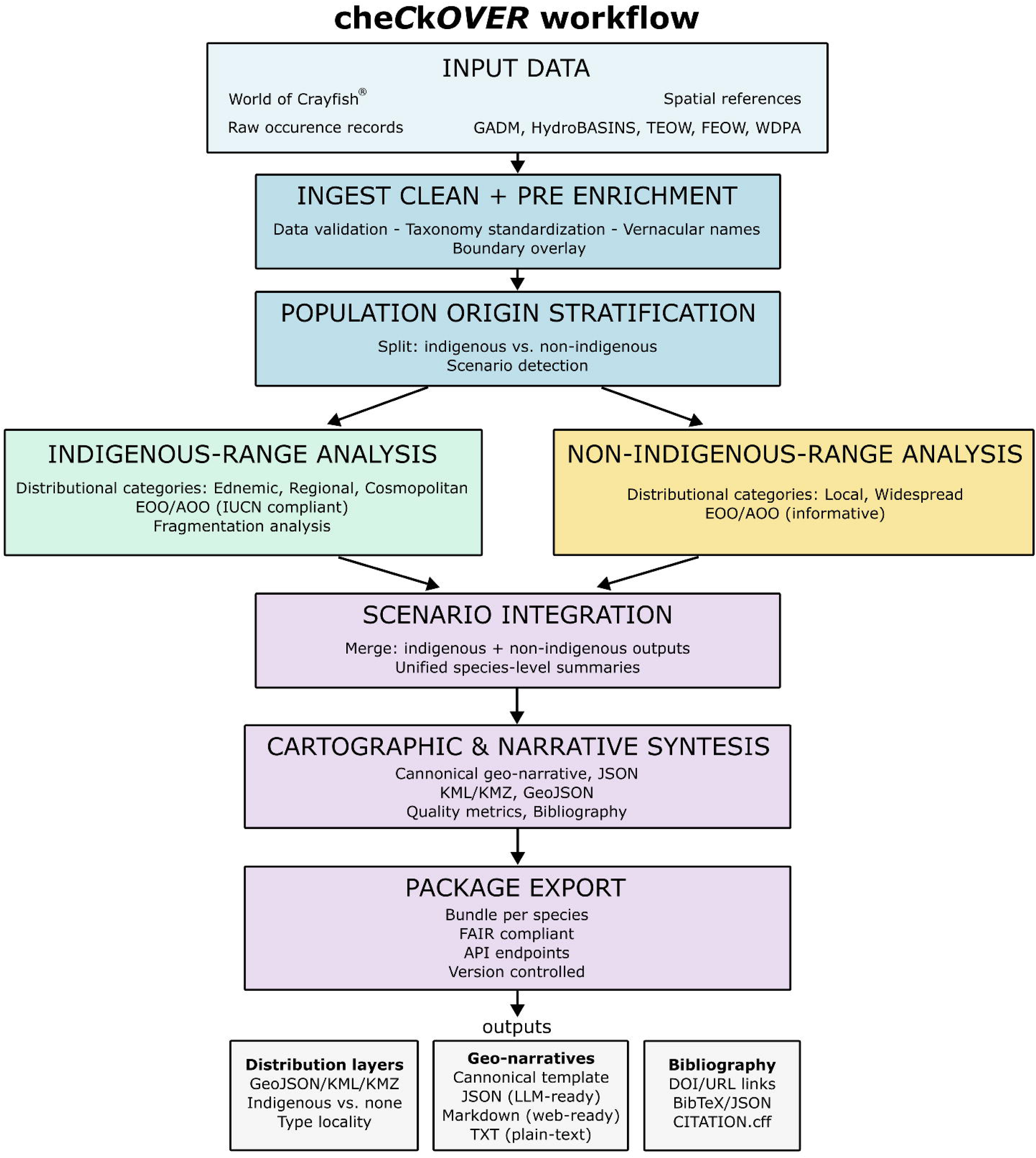
Conceptual workflow of the che*C*k*OVER* framework. The system is organized around five conceptual components: (1) integrated biodiversity input databases; (2) data ingestion, cleaning, and enrichment; (3) analytical processing of distributional and spatial metrics; (4) automated report and narrative generation; and (5) production of modular human- and machine-consumable outputs. Internally, these components are implemented through seven sequential processing phases that stratify species by population origin (indigenous vs. non-indigenous), enabling IUCN-aligned conservation metrics while simultaneously tracking invasion dynamics. The workflow supports both immediate decision-ready reporting and continued context-aware analysis through AI and LLM-based systems, with all outputs embedding provenance metadata for transparent source attribution.

#### Phase 1: Data ingestion, standardization, and pre-enrichment

Raw occurrence records are loaded, cleaned, and standardized (species names, coordinates, status, temporal data). Vernacular names are harvested from authoritative sources and embedded with ISO 639-3 language tags. Initial geospatial enrichment assigns continental context and administrative boundaries (countries, subnational units via GADM v4), establishing the foundational geographic framework for subsequent analyses.

#### Phase 2: Population-origin stratification

To maintain compatibility with IUCN Red List methodology while simultaneously supporting invasion biology workflows, the framework stratifies records by population origin (indigenous vs. non-indigenous). Species are classified into three scenarios: (1) indigenous-only, (2) non-indigenous-only, or (3) combined. This stratification guides all downstream metric calculations and ensures that conservation-relevant indicators (EOO, AOO, fragmentation signals) derive strictly from native-range data, while invasion dynamics are tracked separately.

#### Phase 3: Indigenous-range analysis

For native populations (including type locality), the framework computes distributional categories (endemic, regional, cosmopolitan) based on range extent and geographic scope. Occurrence records are overlaid with standardized geospatial layers (ecoregions, protected areas, hydrographic basins) to generate species-level summaries. For endemic and regional taxa, a conservative fragmentation signal is computed via adaptive spatial clustering of native localities, providing early indications of potential range disjunction without constituting a formal connectivity assessment.

#### Phase 4: Non-indigenous-range analysis

Alien, introduced, or cryptogenic populations are processed through a parallel branch using the same geospatial layers but without fragmentation analysis (as range structure reflects human-mediated dispersal rather than natural processes). Non-indigenous metrics (EOO, AOO, basin occupancy) are computed for informative purposes and support biosecurity prioritization, but do not contribute to conservation status assessments.

#### Phase 5: Scenario integration

For species occurring in both native and introduced contexts (Scenario 3), the framework merges data from both branches into unified outputs that clearly distinguish indigenous from non-indigenous components. This ensures that species pages present complete distributional footprints while preserving the methodological separation required for IUCN-aligned assessments.

#### Phase 6: Narrative and cartographic synthesis

Species-level outputs are generated following a standardized canonical geo-narrative template (Supplementary File S1), covering taxonomic identity, indigenous and non-indigenous ranges, temporal patterns, data quality, and provenance. Narratives are delivered in multiple formats: human-readable text (Markdown, plain text), structured JSON optimized for LLM consumption, and machine-readable metadata files. Geospatial products (KML/KMZ, GeoJSON) visualize indigenous and non-indigenous ranges using distinct color schemes, with type localities highlighted when available.

#### Phase 7: Species package assembly

Final outputs are bundled into self-contained, publication-ready packages, each including maps, narratives, quantitative tables, citation files (BibTeX, JSON, CITATION.cff), and a structured manifest. Privacy protection is ensured by aggregating spatial information to administrative or ecological units, preventing exposure of precise coordinates for sensitive species.

All outputs follow FAIR principles, include output-level versioning (package revision, timestamp, and manifest), and are exposed through documented, API-ready endpoints to support integration with external biodiversity infrastructures and AI-based systems [46]. The framework implements incremental update capability: when new occurrence records are added, only modified species are reprocessed, with results compared against previous versions to track distributional changes over time. This version-aware design ensures computational efficiency for large datasets while maintaining full reproducibility and traceability.

The che*C*k*OVER* pipeline was developed within a modern integrated development environment combining Claude (Anthropic) AI-assisted code generation [47] with expert-driven manual refinement. AI-generated code drafts were iteratively reviewed, validated, optimized, and extended by the authors using real-world occurrence data subsets. This hybrid development workflow ensured reproducibility, code transparency, and compliance with biodiversity informatics best practices [48,49].

### Data inputs and preprocessing

che*C*k*OVER* relies on streamlined occurrence datasets that are easy to maintain and extend over time. The framework was developed and validated using the World of Crayfish^®^ (WoC^®^) database [10], but accepts occurrence data from any taxonomic source following the same core schema. Only core fields required for geo-narrative and quantitative metric generation are used: the valid Latin binomial of the taxon (with synonymy resolved via external APIs), population origin (indigenous vs. alien/introduced), year of observation, geographic coordinates in standardized WGS84 decimal degrees, and a location-accuracy tag. Bibliographic references for each record, including DOI/URL where available, are retained for traceability and citation.

All occurrence records are processed through a series of validation and standardization operations using the tidyverse suite. Species names are cleaned, coordinates coerced to WGS84, population origin normalized (native/alien/introduced/cryptogenic), and accuracy flagged (high/low). Duplicate records (species–year–coordinates–source) are removed, ensuring consistency and reproducibility across datasets. Following preprocessing, records are stratified by population origin, creating two parallel processing branches (indigenous and non-indigenous) while maintaining a unified master dataset for scenario detection. Species are then classified into three scenarios based on their population-origin profiles: Scenario 1 (indigenous-only), Scenario 2 (non-indigenous-only), or Scenario 3 (both population types present). This classification guides downstream spatial resolution choices (e.g., finer HydroBASINS levels for endemic taxa, coarser resolutions for cosmopolitan taxa) and ensures internal consistency across quantitative metrics and visual outputs.

To support multilingual, real-world queries, vernacular (common) names are not required as inputs; instead, they are harvested downstream in a dedicated step from authoritative and community-curated services (e.g., ITIS; https://www.itis.gov/), or supplemented with expert-reviewed multilingual names compiled in a static lookup table (Table S2). This step is implemented using the *ritis* package for querying authoritative vernacular-name services. All vernacular names are normalized with ISO 639-3 language tags. The resulting language-aware name lists are embedded in species outputs to improve discoverability without complicating the primary input schema.

### Geospatial enrichment

To generate narrative-ready outputs, che*C*k*OVER* links validated occurrence records to a suite of standardized geospatial layers through a two-stage enrichment process. Stage 1 (pre-stratification) assigns continental context and administrative boundaries to all records prior to population-origin splitting, establishing a unified geographic framework. Stage 2 (post-stratification) performs branch-specific spatial overlays with ecoregions, protected areas, and hydrographic basins, with resolution and layer selection adapted to each population type’s biogeographic context.

These layers provide named, interpretable geographic units—countries, administrative regions, river basins, ecoregions, and protected areas—that allow species distributions to be described using stable, widely recognized terminology. Ensuring correct names for these units is essential, and the framework therefore integrates both programmatic attribution (via R packages) and reproducible lookup tables for cases where names are absent from the original polygons.

#### Stage 1: Universal geographic context

Country-level boundaries are sourced from Natural Earth (via the *rnaturalearth* package), while sub-national administrative units are retrieved through the geodata package, which provides standardized access to GADM v4. Continental assignments are derived from Natural Earth medium-scale vector data. These layers are applied uniformly to all occurrence records regardless of population origin, ensuring that baseline geographic descriptors remain consistent across indigenous and non-indigenous subsets.

#### Stage 2: Branch-specific biogeographic enrichment

Following population-origin stratification, indigenous and non-indigenous records are processed through separate but parallel spatial overlay workflows. This separation ensures that IUCN-aligned conservation metrics derive exclusively from native-range data while enabling comprehensive tracking of introduced populations for invasion biology applications.

Freshwater contexts are captured through HydroBASINS hierarchical watershed boundaries (levels L06–L10) and HydroATLAS hydro-environmental descriptors, while WWF Terrestrial Ecoregions of the World (TEOW) and Freshwater Ecoregions of the World (FEOW) are assigned using the *ecoregions* and *feowR* packages. Because HydroBASINS polygons do not include hydronym attributes, basin names were derived through a dedicated offline geoprocessing step. HydroBASINS units (L06, L08, L10) were spatially intersected with river-line datasets from the Global Runoff Data Centre (for L06–L08) and high-resolution OpenStreetMap extracts (for L10). Each basin polygon was assigned the name of the longest or most representative river crossing its area. This workflow yielded a static hydronym lookup table linking HydroBASINS IDs to river names (Table S3), now integrated into the che*C*k*OVER* pipeline. When multiple HydroBASINS IDs share the same river name (e.g., multiple segments of the Danube), records are aggregated by basin name and summed for reporting, ensuring that outputs present deduplicated, human-readable geographic units rather than technical polygon identifiers. In contrast, TEOW polygons include standardized ecoregion names directly via the ecoregions package; therefore, no additional processing was required. FEOW polygons, however, lack embedded name attributes in some distributions, and a reproducible lookup table linking FEOW IDs to ecoregion names was compiled (Table S4). Protected areas were intersected using the World Database on Protected Areas (WDPA, Protected Planet), retrieved programmatically via the *wdpar* package to ensure license-compliant access.

For non-indigenous populations fragmentation analysis is omitted, as introduced-range structure reflects human-mediated dispersal patterns rather than natural biogeographic processes. This design ensures that biosecurity-relevant spatial summaries (basin occupancy, protected-area overlap) are generated without conflating anthropogenic range expansion with evolutionarily meaningful fragmentation signals.

All geospatial operations—cropping, overlays, joins, reprojections, and geometry handling—are executed using the *sf* package, which provides high-performance spatial data processing and consistent handling of modern GIS geometries.

### Range metrics and distributional categorization

For range metrics, the framework computes extent of occurrence (EOO) for native ranges as minimum convex polygons encompassing native validated records, following IUCN guidelines [50]. To avoid artificial inflation in species with disjunct ranges across continents or islands, EOO is calculated as the sum of convex hulls per contiguous landmass (or freshwater system), preventing polygons spanning oceans. Area of occupancy (AOO) is calculated as the number of occupied grid cells (typically 2×2 km) multiplied by cell area. Occurrences flagged as extinct are removed prior to spatial overlays and range calculations, ensuring that EOO and AOO represent only the current distribution. Extinct localities remain available for reporting but do not influence geospatial metrics. All outputs distinguish between native and alien ranges to maintain ecological relevance and ensure compatibility with IUCN conservation standards. To maintain full compatibility with IUCN Red List methodology, alien/introduced occurrences are never included in metric calculations, but are retained for spatial reporting and visualization. Accordingly, all geospatial display products (GeoJSON, KML/KMZ) show both native and alien ranges when present, while the corresponding EOO/AOO values included in reports and JSON summaries derive strictly from the native subset of records.

To contextualize the breadth of native distributions prior to spatial enrichment, che*C*k*OVER* classifies taxa into three distributional categories using native-range occurrence statistics only. (i) Endemic taxa are defined as species with a native extent of occurrence (EOO) ≤5,000 km², restricted to ≤2 countries within a single continent. These thresholds are consistent with restricted-range criteria commonly applied to freshwater invertebrates [51] and reflect the naturally fragmented distributions of Astacidea across isolated drainage systems [52,53]. (ii) Regional taxa are species whose native ranges exceed endemic thresholds but remain confined to a single continent, regardless of any secondary introductions outside their native area. (iii) Cosmopolitan taxa are defined as species whose native ranges span more than one continent, indicating broad historical biogeographic distributions rather than anthropogenic spread.

Records outside the native range (alien, introduced, or cryptogenic occurrences) are explicitly retained and analyzed separately for descriptive statistics, protected-area intersections, basin-level summaries, and cartographic outputs, but they do not influence native-range classification. This separation ensures that biogeographic categorization reflects evolutionary and historical range structure, while still capturing the full contemporary distributional footprint of each species. This native-range classification guides downstream spatial resolution choices (e.g., finer HydroBASINS levels for endemic taxa and coarser resolutions for cosmopolitan taxa), ensuring internal consistency across quantitative metrics and visual outputs.

### Fragmentation analysis (indigenous populations only)

For endemic and regional taxa, che*C*k*OVER* additionally computes a simple distributional fragmentation signal to summarize the spatial cohesion of known native localities. The analysis is performed on straight-line (Euclidean) distances, calculated after transforming occurrence coordinates to an equal-area projection (EPSG:6933) to ensure metric accuracy. A full pairwise distance matrix is generated, and the mean inter-point distance is used as an adaptive species-specific threshold for hierarchical clustering (complete linkage). This approach allows fragmentation detection to scale naturally with each species’ spatial configuration. Clusters are derived using *hclust()* and *cutree()* at the adaptive height. Species with fewer than five valid locality records, as well as cosmopolitan taxa whose ranges are inherently multi-continental, are excluded from clustering. Non-indigenous populations are excluded from fragmentation analysis by design, as introduced-range structure reflects human-mediated dispersal rather than natural connectivity patterns.

### Computational optimization

To minimize computational load during large-scale intersection operations, che*C*k*OVER* implements an optimized, multi-stage spatial overlay procedure. For each species, the framework first computes an early EOO polygon, buffered to account for spatial uncertainty. All reference layers (countries, basins, protected areas) are dynamically cropped to this bounding region prior to point-level intersections, reducing redundant geometry checks. Spatial joins are executed in batches using indexed geometries (*sf* + *S2* engine), followed by tabular aggregation per species and per geographic unit. When applicable, HydroBASINS levels are dynamically selected (L10 for endemic, L8 for regional, L6 for cosmopolitan taxa), and protected areas are pre-filtered to remove marine or micro-polygons. This adaptive workflow reduces memory requirements and computation time while maintaining full spatial accuracy.

### Narrative generation and outputs

The core function of che*C*k*OVER* is to transform enriched species occurrence records into structured geo-narratives that are both human-readable and machine-interoperable. Species-level outputs follow a standardized canonical geo-narrative template (Supplementary File S1), implemented using the glue package, ensuring consistency across taxa and enabling fully reproducible reporting for both human users and automated systems.

#### Canonical template structure

The canonical template organizes biodiversity knowledge into five standardized sections: (1) Taxonomic Identity, including scientific name, higher taxonomy (Order > Superfamily > Family), vernacular names with full language names (not ISO codes in output text), and type locality when available; (2) Indigenous Range Overview, covering distribution category (endemic/regional/cosmopolitan), EOO/AOO metrics, geographic distribution (countries, subnational units, hydrographic basins—all sorted by record count descending), protected area coverage, biogeographic context (TEOW/FEOW ecoregions), temporal context, auto-generated contextual statements, range dynamics (if extinctions documented), and fragmentation assessment (endemic/regional only); (3) Non-Indigenous Range Overview (informative section for species with introduced populations), reporting EOO/AOO, countries, basins, and temporal patterns separately from native metrics; (4) Data Quality, Traceability, and Provenance, documenting total records, indigenous/non-indigenous breakdown, high-accuracy records (coordinate uncertainty <1 km), data quality score (0–100 composite), bibliographic coverage (total references vs. DOI-linked references), raw data provenance (WoC^®^ citation), processing framework provenance (che*C*k*OVER* version, processing date, data snapshot date), and interpretation note (disclaimer about non-IUCN status); and (5) Formal Narrative Summary, a 300–500 word human-readable synthesis structured in five paragraphs covering taxonomic identity, indigenous range geography/biogeography, conservation context (protected areas, fragmentation, extinctions), non-indigenous populations (if applicable), and data quality/provenance.

Validated data are organized into predefined narrative blocks following this template structure. While the framework relies on a fixed narrative structure to guarantee comparability and reproducibility, it remains extensible, allowing the addition of new sections or user-defined analytical queries as the system evolves.

For Scenario 1 species (indigenous-only), outputs derive entirely from the indigenous processing branch. For Scenario 2 species (non-indigenous-only), Section 2 (Indigenous Range) is omitted, and Section 3 (Non-Indigenous Range) becomes the primary distributional summary. For Scenario 3 species (both population types), the framework merges data from both branches: Section 2 presents native-range metrics, Section 3 presents introduced-range metrics, and both are clearly demarcated to prevent conflation of native vs. alien distributions in downstream analyses.

Predefined query blocks are embedded within each narrative to support automated responses to frequently asked questions, while the structured JSON outputs can be directly consumed by large language models (LLMs) to address free-form prompts. When available, type localities are highlighted as key descriptive attributes, explicitly linked to their bibliographic sources, and incorporated into the indigenous range narrative. Vernacular names harvested during earlier processing steps are automatically embedded in the JSON narratives using ISO 639-3 language tags internally, but are rendered with full language names (e.g., “English: noble crayfish” not “eng: noble crayfish”) in human-readable outputs, enabling accurate querying with local or non-scientific terminology without compromising taxonomic precision.

Each narrative includes a concise distributional summary, reported both in JSON and tabular formats, presenting key metrics such as EOO, AOO, total numbers of validated occurrences, and counts of records annotated as extinct. For endemic and regional taxa, che*C*k*OVER* additionally reports a conservative distributional fragmentation signal derived exclusively from indigenous occurrence records. This signal contextualizes the spatial cohesion of known localities by identifying geographically isolated clusters using adaptive, species-specific clustering thresholds. Fragmentation outputs include (i) the number of detected spatial clusters, (ii) their relative sizes expressed as occurrence counts and percentages, and (iii) a categorical indicator of whether fragmentation was detected (“not detected” for a single cohesive cluster; “detected” when multiple clusters are present). These signals are intended as supportive descriptors aligned with IUCN criteria B and D [50], without constituting a formal Red List assessment.

Final outputs are delivered in multiple interoperable formats: (i) canonical narratives rendered via *rmarkdown* into Markdown files containing all five template sections; (ii) formal narrative summaries (Section 5 only) exported as both plain text (TXT) and structured JSON for streamlined LLM consumption; (iii) structured JSON files generated using *jsonlite*, optimized for programmatic access and LLM-based workflows, with embedded metadata blocks for each section; and (iv) geospatial artefacts (KML/KMZ and GeoJSON) supporting visualization and spatial integration. Each JSON narrative embeds a persistent pointer to a de-duplicated, species-level bibliography, ensuring full provenance, traceability, and unambiguous citation even when accessed independently. A permanent “How to cite this resource” entry is maintained in the che*C*k*OVER* documentation and will be updated following publication with the definitive citation and DOI.

### Implementation

The che*C*k*OVER* initial scripts were drafted and subsequently validated, optimized, and extended by the authors, ensuring reproducibility and compliance. che*C*k*OVER* was implemented in R v.4.3.3 [54] as a modular package, with all functionality exposed through callable functions (Table 1). The workflow integrates sequential modules for ingestion, enrichment, reporting, and packaging, yet each component can also be executed independently. For transparency and reproducibility, all raw scripts are distributed within the package (*inst*/*scripts*_*raw*/), enabling stepwise execution when required. The pipeline is orchestrated by the *run_pipeline()* function, which reads a central configuration file (YAML), sets paths and column mappings, filters taxa, and executes modules sequentially or selectively with full logging and error handling.

**Table 1.** Core modules of the che*C*k*OVER* R package and their primary functions. A full inventory of supporting R packages required by the pipeline, together with their roles and reference URLs, is provided in Table S5.

| Module | Core function |
| --- | --- |
| *ingest_clean()* | Ingests raw occurrence records (XLSX/TSV/DB), validates the input schema through configurable column mapping (default WoC<sup>®</sup>-compatible), and standardizes core fields. Species names are cleaned, coordinates coerced to WGS84, population origin normalized (native/type locality/alien/introduced/cryptogenic), and accuracy flagged. Duplicate records (species–year–coordinates–source) are removed; taxonomy may optionally be resolved via external APIs. Exports standardized tabular (CSV/TSV/RDS) and spatial (*sf*) objects. |
| *vernacular_names()* | Retrieves vernacular (common) names from prioritized sources (e.g., |
|  | ITIS), normalizes spelling, removes duplicates, applies ISO 639-3 tags internally, and supports manual curation. Outputs consolidated name tables (TSV) and injects them into JSON narratives with full language names (e.g., "English: noble crayfish") to enable queries using non-scientific names. |
| *split_by_population()* | Stratifies occurrence records by population origin (indigenous vs. non-indigenous), creating two parallel datasets while maintaining a unified master dataset. Enables IUCN-aligned metrics from native records while simultaneously tracking invasion dynamics. Exports separate TSV files for each population type. |
| *detect_species_scenarios()* | Classifies species into three scenarios based on population-origin profiles: Scenario 1 (indigenous-only), Scenario 2 (non-indigenous-only), or Scenario 3 (both population types present). This classification guides downstream processing branches and output merging strategies. Exports scenario classification table (TSV/JSON). |
| *enrich_with_continents()* | Assigns continental context to all occurrence records using Natural Earth medium-scale vector data. Applied uniformly before population-origin split to establish baseline geographic framework. |
| *enrich_with_gadm()* | Overlays occurrence records with country and subnational administrative boundaries (GADM v4.1) using the *geodata* package. Applied before population split to ensure consistent political geography across both branches. |
| *calculate_indigenous_metrics()* | Computes distributional categories (endemic/regional/cosmopolitan), EOO, AOO, and preliminary range statistics exclusively from native occurrence records. Implements IUCN-aligned thresholds (endemic: $EOO \leq 5,000 \text{ km}^2$, $\leq 2$ countries, single continent). Guides downstream spatial resolution choices (HydroBASINS level selection). |
| *analyze_fragmentation()* | Detects distributional fragmentation patterns in endemic and regional taxa via adaptive Euclidean-distance clustering. Generates pairwise distance matrix, uses mean distance as species-specific threshold, and applies hierarchical clustering to identify spatially disjunct groups. Outputs cluster count, sizes (counts and percentages), and categorical fragmentation flag. Applied only to indigenous populations; cosmopolitan and non-indigenous taxa excluded by design. |
| *enrich_indigenous_spatial()* | Overlays native occurrence records with biogeographic layers: hydrographic basins (HydroBASINS L06/L08/L10, resolution adaptive to distributional category), ecoregions (FEOW/TEOW), and protected areas (WDPA). Implements optimized workflow: early EOO polygon crops reference layers, indexed geometry operations (*sf* + *S2* engine) batch-process intersections. Basin names assigned via offline hydronym lookup (Table S3); FEOW names via lookup table (Table S4). Aggregates records by basin name (deduplicating multiple polygon IDs for same river). Outputs enriched per-record spatial attributes and summary statistics. |
| *generate_indigenous_reports()* | Compiles quantitative species reports for native populations: record totals, temporal coverage, geographic tallies (countries, subnational units, basins, protected areas—all sorted by record count descending), density metrics, range statistics (EOO, AOO), fragmentation signals, and extinction summaries. Exports tidy tables (TSV), structured JSON summaries, and markdown reports. |
| *calculate_non_indigenous_metrics()* | Computes distributional categories (local/widespread), EOO, AOO, and range statistics exclusively from alien/introduced occurrence records. Metrics are informative (not IUCN-aligned) and support biosecurity prioritization. No fragmentation analysis applied. |
| *enrich_non_indigenous_spatial()* | Overlays alien/introduced records with the same geospatial layers as indigenous branch (basins, ecoregions, protected areas) but omits |
|  | fragmentation analysis. Ensures biosecurity-relevant spatial summaries without conflating human-mediated dispersal with natural biogeographic patterns. |
| *generate_non_indigenous_reports()* | Compiles quantitative reports for introduced populations: record totals, temporal coverage, geographic tallies (countries, basins, protected areas), range statistics. Statistics reported separately from native metrics to maintain IUCN compatibility. Exports TSV, JSON, and markdown. |
| *merge_scenario3_reports()* | For Scenario 3 species (both population types), merges data from indigenous and non-indigenous branches into unified outputs that clearly distinguish native from alien components. Ensures species pages present complete distributional footprints while preserving methodological separation required for IUCN assessments. |
| *generate_canonical_narratives()* | Generates structured geo-narratives following the canonical template (Supplementary File S1). Narratives cover five sections: (1) Taxonomic Identity (scientific name, higher taxonomy, vernacular names, type locality), (2) Indigenous Range Overview (distribution category, EOO/AOO, countries, subnational units, basins, protected areas, ecoregions, temporal context, fragmentation), (3) Non-Indigenous Range (if applicable), (4) Data Quality and Provenance (record counts, accuracy metrics, bibliographic coverage, processing metadata), (5) Formal Narrative Summary (300–500 word human-readable synthesis). Outputs canonical markdown (all sections), formal narrative TXT (Section 5), and formal narrative JSON (Section 5 structured). |
| *generate_maps()* | Produces standardized geospatial layers (KML/KMZ, GeoJSON) using the sf framework. Generates three map types per species: EOO convex hulls (yellow polygons), AOO grid cells (2×2 km, yellow), and HydroBASINS occupancy with population-type color coding (native: orange, introduced: purple, mixed: red-orange for Scenario 3). Type localities marked with red 2×2 km squares when available. Map resolution adapts to distributional category (L10 endemic, L8 regional, L6 cosmopolitan). Privacy-protected via spatial aggregation. |
| *generate_all_citations()* | Aggregates and de-duplicates bibliographic references from occurrence records, preserves DOIs/URLs, and generates multiple citation formats: APA-formatted lists, BibTeX files, structured JSON, and CITATION.cff metadata files. Embeds provenance links in all JSON outputs to enable source attribution in LLM-mediated workflows. |
| *export_species_packages()* | Bundles all species-level outputs into distributable packages: geospatial layers (KML/KMZ, GeoJSON), canonical narratives (Markdown, TXT, JSON), quantitative tables (CSV, TSV), citations (BibTeX, JSON, CFF), and metadata files. Generates manifests, checksums, and versioned outputs to support reproducibility and longitudinal tracking. Each package includes README documentation. |
| *run_pipeline()* | Orchestrates sequential or selective module execution across all seven processing phases. Supports YAML config files, CLI parameters, logging, resume mode (detects existing outputs, skips completed modules), and reproducibility reporting. Records version history and reports differences between current and earlier processed outputs. Implements optional parallel execution for indigenous/non-indigenous branches (Linux/macOS via multicore, Windows via multisession). |

### Deployment architecture and API accessibility

che*C*k*OVER* outputs clear, well-structured, machine-consumable metadata that allow seamless integration in complex processing workflows and support efficient integration with existing LLM services through RESTful API endpoints (e.g., /api/llm/{species_name}). The framework supports both local execution and server deployment configurations, ensuring applicability across institutional scales from individual researchers to large biodiversity platforms.

We comply with all the important aspects for providing a solution that is both API-ready and LLM-ready: (1) machine consumable data and metadata, through standardized formats (GeoJSON, KML/KMZ, MD, TSV etc.); (2) rich error semantics that improve API discoverability and help AI agents and API integrators to understand data structure and requirements, behavior and recover from errors; (3) speed and reliability ensured by a containerization-layered technical stack, which provides seamless horizontal scalability at each level: repository, processing, data conversions; (4) consistent REST end-points design which, together with the standardized data and querying formats provides predictability; (5) complete and well structured documentation available through OpenAPI specification.

Data storage combines species-level file hierarchies with a MariaDB relational database that indexes taxonomic, geographic, and metadata attributes for efficient querying and retrieval. For production environments, species packages can be exposed via documented API endpoints that return structured JSON responses optimized for retrieval-augmented generation workflows. The system is fully containerized (Docker/Singularity), supporting horizontal scalability at each processing layer and ensuring portability across computing environments.

### Reproducibility and privacy protection

All code developed for che*C*k*OVER* is released as open source and publicly available in a version-controlled *GitHub* repository. To ensure portability across systems, che*C*k*OVER* is fully containerized and can be deployed both locally and within Docker or Singularity environments. Stable releases are versioned and archived, ensuring long-term reproducibility. This modular and container-ready design facilitates straightforward maintenance, interoperability with other biodiversity infrastructures, and adaptability across taxonomic groups.

Sensitive species records require protection against potential misuse, such as illegal harvesting or habitat disturbance. che*C*k*OVER* ensures that raw geographic coordinates are never released in its outputs. Instead, all products—including dynamic maps, quantitative summaries, and biogeographic narratives—are based on aggregated or generalized spatial information. Geographic descriptors are expressed in terms of administrative or ecological units (e.g., countries, basins, ecoregions, protected areas), while statistical summaries report counts, densities, and extents without exposing point-level data. To ensure full Unicode compatibility across platforms (Windows/macOS/Linux), all tabular outputs are exported in TSV format, which avoids encoding corruption of special characters and remains interoperable with biodiversity infrastructures.

## Results

### Implementation and dataset processing

We implemented che*C*k*OVER* using global crayfish (Astacidea) occurrence records from the WoC^®^ database [10], comprising 115,486 occurrence records for 465 taxa spanning 1758–2025, contributed by the World of Crayfish^®^ consortium members (contributor details provided in Table S1). While designed as a taxon-agnostic framework, the WoC^®^ dataset provided comprehensive validation due to its global scope and well-documented metadata.

The data ingestion and preprocessing module successfully processed 96.1% of input records (111,729 from 116,274 raw records), with 3,801 records removed due to coordinate validation failures and 744 duplicates eliminated. Taxonomy resolution through external APIs standardized 465 species names, while vernacular name harvesting from ITIS and expert inputs yielded 713 common names across 35 languages. Multiple language versions of recognized common names were retained using special characters without forcing translation into English. Type locality information was successfully identified and preserved for 120 species (25.8% of total), each linked to original taxonomic descriptions with full bibliographic references.

Geospatial enrichment successfully assigned all validated records to standardized geographic units spanning 97 countries, hierarchically nested hydrographic basins (HydroBASINS levels L06–L10), and terrestrial and freshwater ecoregions (TEOW/FEOW), providing comprehensive geographic coverage for demonstrating framework scalability. Protected area coverage was calculated for all species distributions using WDPA-designated boundaries, enabling assessment of conservation context and identification of protection gaps.

### Quantitative descriptors and metrics

All quantitative metrics used for conservation assessment—including EOO, AOO, distributional categories (endemic, regional, cosmopolitan), and fragmentation signals—were calculated exclusively from native occurrence records, ensuring compatibility with IUCN Red List methodology. Spatial statistics derived from geographic overlays (country counts, basin tallies, ecoregion assignments, protected area intersections) are computed for both native and alien records but reported separately in all outputs.

Although both the IUCN Red List of Threatened Species (RLTS) and the Red List of Ecosystems (RLE) apply structured risk assessment frameworks, they address fundamentally different conservation targets and are not interchangeable. RLTS evaluates extinction risk at the species level, where evidence of population reduction is central, whereas RLE focuses on ecosystem collapse inferred from changes in distribution, function, and integrity [50,55]. Consequently, inference rules and criteria developed for ecosystem-level assessments cannot be directly transposed to species-level Red List evaluations. Within che*C*k*OVER*, distributional metrics such as EOO and AOO are therefore treated strictly as descriptive spatial evidence supporting range delineation, spatial structure, and temporal comparison, and not as direct indicators of threat or decline under RLTS Criterion A, where pressures are only relevant insofar as they can be linked to demonstrable population reduction.

che*C*k*OVER* framework computes a standardized set of quantitative distributional descriptors from both indigenous (Table 2) and non-indigenous (Table 3) occurrence records, while maintaining strict stratification for reporting and visualization purposes. For native-range assessments, species were classified into three distributional categories based on indigenous range extent and geographic scope: 171 endemic taxa (36.8%), 287 regional taxa (61.7%), and 5 cosmopolitan taxa (1.1%). These classifications guide IUCN-aligned conservation metrics and adaptive spatial resolution assignment.

**Table 2.**
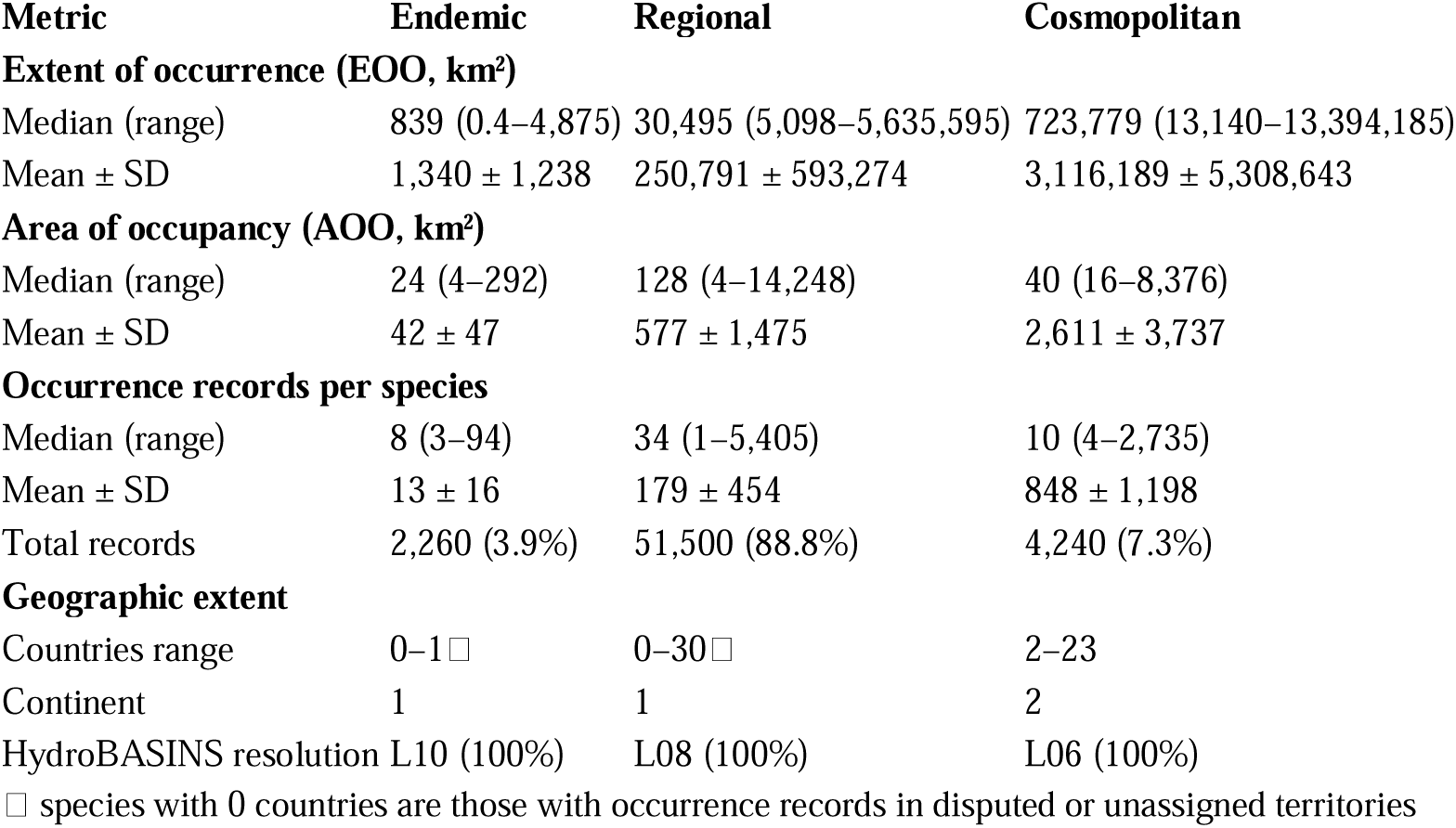
Distributional metrics by biogeographic category for crayfish species with native-range occurrences. Species classified as Endemic (n=171), Regional (n=287), or Cosmopolitan (n=5). All metrics calculated from indigenous records only (58,000 total). Values shown as median (range). EOO = extent of occurrence; AOO = area of occupancy.

**Table 3.**
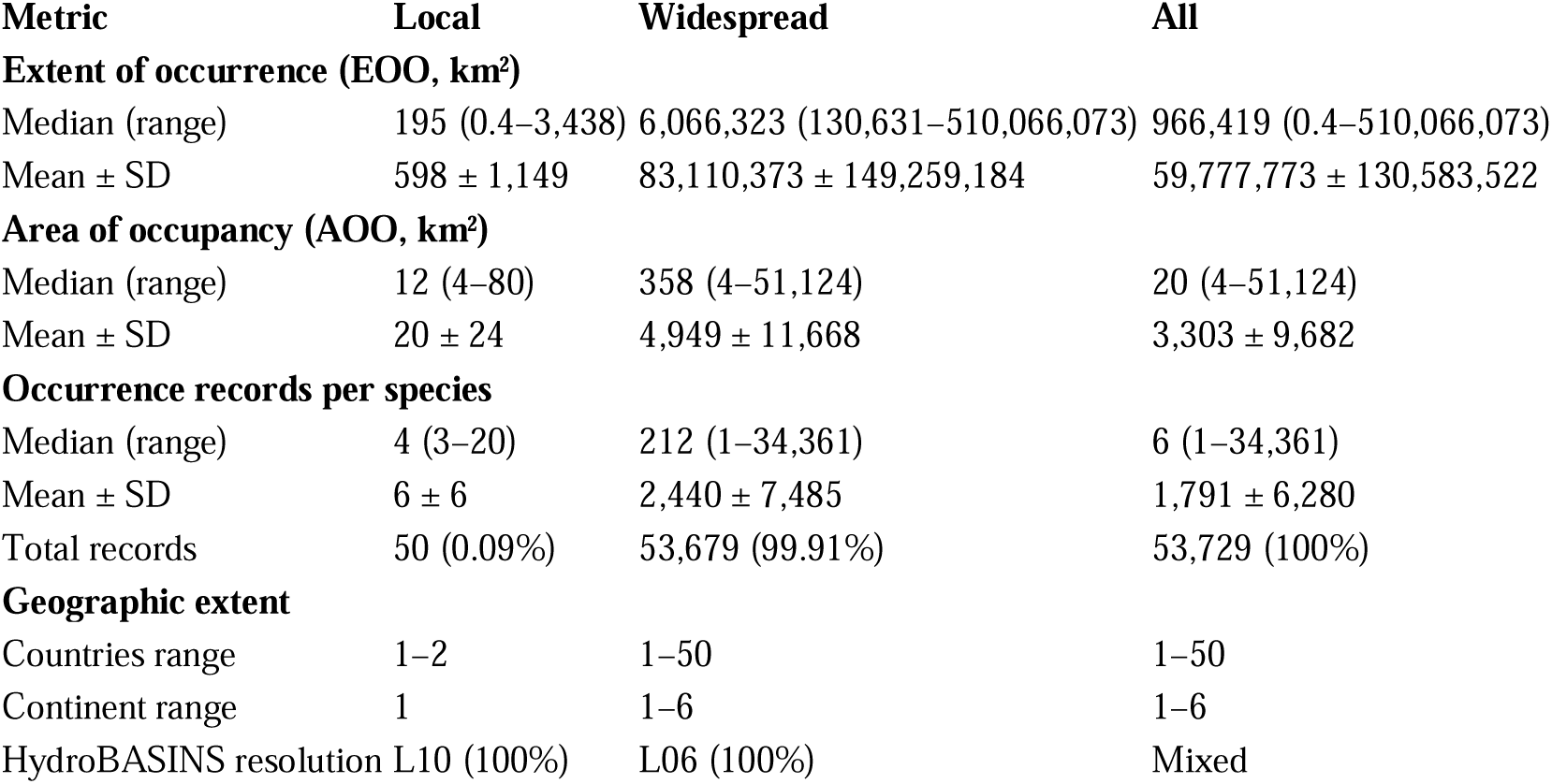
Non-indigenous range metrics for invasive crayfish species. Species classified as Local (n=8) or Widespread (n=22) based on introduced-range extent. All metrics from non-indigenous records only (53,729 total). Values shown as median (range). These metrics are informative only and do not contribute to conservation assessments. EOO = extent of occurrence; AOO = area of occupancy.

For invasion biology applications, non-indigenous ranges were analyzed separately across 30 species documented with alien populations (Table 3). Range extent varied dramatically, with 8 species exhibiting localized distributions (median 4 alien records, restricted to 1–2 countries) and 22 species displaying widespread invasions (median 212 alien records, spanning multiple countries or continents). The global invasion footprint is dominated by a small number of highly successful invaders: *Procambarus clarkii* (34,361 alien records across 50 countries on 5 continents), *Faxonius limosus* (7,484 records, 23 countries), and *Pacifastacus leniusculus* (6,408 records, 29 countries spanning 3 continents) collectively account for 89.8% of all non-indigenous occurrence data. Despite representing only 6.5% of all processed species (30 of 465), these invasive populations generated 53,729 alien occurrence records—approaching parity with the 58,000 indigenous records from 463 native-range species—reflecting a disproportionate occurrence indexing effort.

The distributional analysis revealed significant variation in species’ geographic extents. Endemic species showed restricted EOO values (median: 839 km²), while cosmopolitan taxa displayed extensive ranges spanning multiple continents. Alien range stratification identified 30 species with documented non-indigenous records, representing 6.5% of the processed dataset. For endemic and regional taxa, che*C*k*OVER* assessed the spatial cohesion of native occurrence records through a simple fragmentation analysis. Of 458 species in the endemic and regional categories, 362 (79.0%) had sufficient occurrence data (≥5 validated indigenous localities) for reliable clustering analysis. Distributional fragmentation was detected in all 362 analyzed species, with cluster counts ranging from 2 to 25 (median: 5 clusters), reflecting varying degrees of spatial disjunction across native ranges. The remaining 96 species (21.0%) had insufficient data points for clustering. Cosmopolitan species were excluded from this analysis by design.

### Output generation and validation

The framework successfully generated complete species packages for all 465 processed taxa. Each package contains standardized outputs across multiple formats, including visual representations (displaying native and alien ranges using distinct colors, with type locality when available), geo-narratives (TXT, MD, JSON), and bibliographic metadata provided both as APA-formatted citations and machine-readable CITATION.cff. Geospatial outputs were generated at resolutions scaled to distributional categories: endemic species were exported using high-resolution basin units (HydroBASINS L10), regional species used intermediate resolution (L08), and cosmopolitan species used broader basin levels (L06). Native and alien ranges are visually distinguished using distinct colors in all map outputs. All exported layers incorporate privacy protection by aggregating spatial information instead of exposing raw coordinate data.

Geo-narrative generation produced structured JSON outputs optimized for LLM consumption alongside human-readable text summaries. Each narrative included standardized sections covering geographic distribution, ecological status, temporal patterns, and data quality assessments. Pre-computed answers to common queries (e.g., “Is this species invasive?”, “What is its biogeographic and preliminary conservation status?”) were embedded to facilitate automated responses.

### Framework performance and scalability

che*C*k*OVER* demonstrated robust performance across the complete crayfish dataset. Processing 465 species with 111,729 validated occurrence records required approximately 41 hours on standard hardware, with linear scaling observed for larger datasets. Peak memory requirements reached approximately 220 GB during intensive geospatial operations, particularly during high-resolution GeoJSON/KML generation and protected-area intersections for species with very large occurrence datasets. The modular architecture enabled selective execution of pipeline components, supporting both complete workflow runs and targeted updates. Individual modules executed independently without framework dependencies, facilitating maintenance and customization for different taxonomic groups or data sources. Output validation confirmed metadata completeness (100% for core descriptors), comprehensive bibliographic traceability with documented provenance chains, and API accessibility supporting programmatic integration with external platforms.

che*C*k*OVER* has modest baseline computational requirements for standard workflows, but resource usage scales with dataset size and the complexity of spatial operations. For small to moderate datasets (e.g., up to a few thousand occurrence records per species), the pipeline typically requires ≤8 GB RAM. Intermediate steps involving spatial overlays (e.g., basin and protected-area intersections) may require in the order of tens of gigabytes of memory. For species with very large occurrence datasets (tens of thousands of records), memory-intensive operations such as high-resolution GeoJSON/KML generation and range polygon rendering (AOO/EOO) can require substantially higher RAM (≥128 GB), particularly when processed at fine spatial resolutions. Disk requirements scale primarily with the number of generated geospatial artifacts and cached intermediate products; several tens to hundreds of gigabytes may be required for full multi-species runs. Users are therefore encouraged to adjust computational resources according to dataset size and intended output resolution.

## Discussion

The crayfish dataset serves as a proof-of-concept demonstration of che*C*k*OVER*’s technical capabilities across diverse distributional scenarios. The quantitative patterns reported reflect the characteristics of available occurrence data rather than definitive biogeographic or ecological assessments. Occurrence datasets inherently reflect reporting effort, accessibility, taxonomic expertise, and monitoring priorities rather than complete sampling of actual distributions. che*C*k*OVER* outputs are designed as standardized, reproducible syntheses of available data to support—but not replace—expert-driven assessments, field validation, and integration with additional evidence sources. The framework’s value lies in its ability to transform heterogeneous occurrence records into consistent, interoperable knowledge products regardless of data quality or taxonomic group, rather than in biological conclusions derived from any single dataset. The following discussion focuses on the framework’s technical design, capabilities, and broader implications for biodiversity informatics.

### User interaction paradigms and accessibility

che*C*k*OVER* is designed to serve diverse user communities through complementary modes of interaction. Direct access via APIs and downloadable species packages supports traditional workflows for researchers, conservation practitioners, and institutions, providing complete datasets with full methodological transparency. In parallel, indirect access through LLM intermediaries could create conversational interfaces, allowing users to query species knowledge without prior taxonomic expertise or database literacy.

A distinctive feature is the integration of multilingual vernacular names, systematically tagged by language. Vernacular names typically reflect local cultural perspectives rather than direct translation of scientific nomenclature. Incorporating them expands accessibility for global users, enabling queries phrased in national languages or traditional ecological terminology to be accurately matched to validated species records. For Indigenous and local communities, whose ecological knowledge is encoded in vernacular systems rather than formal taxonomies, this creates new avenues for recognition and connection. By embedding vernacular names while preserving scientific rigor, che*C*k*OVER* establishes a two-way bridge between cultural knowledge and biodiversity infrastructures.

Beyond technical performance, che*C*k*OVER* aligns with broader principles in interdisciplinary biodiversity research, which emphasize that effective integration requires coordinated attention to people, process, and perspectives within collaborative teams [56]. By combining computational infrastructure with expert validation, multilingual access pathways, and culturally grounded vernacular systems, the framework operationalizes these principles in practice. Its API-ready, AI-streamable JSON outputs and dynamic versioned database facilitate dialogue between data scientists, taxonomists, conservation practitioners, and users rooted in local ecological knowledge, reinforcing the view that durable biodiversity infrastructures emerge not only from technical design, but from sustained mutual respect and shared epistemic perspectives.

### Toward dynamic biodiversity knowledge systems

By embedding API readiness and automated output versioning, che*C*k*OVER* transforms biodiversity repositories from static archives into dynamic knowledge systems. Outputs can be regenerated as databases expand, taxonomies change, or new spatial references become available. This capacity ensures that users—and AI systems—access current information while retaining archived output versions for temporal or methodological analyses.

The framework’s handling of extinction events exemplifies this temporal dimension: documented population losses are preserved as historical context while being excluded from current range metrics, enabling transparent tracking of biodiversity change over time. Such dynamism is particularly valuable in fast-moving contexts such as invasion biology, disease ecology, and climate-change impact assessments [57,58]. The continuous-integration model positions biodiversity infrastructures as living systems—responsive to scientific progress while preserving methodological rigor.

### Transforming biodiversity informatics for the AI era

Digital technologies are becoming indispensable for accelerating biodiversity knowledge and conservation, while also highlighting practical constraints such as accessibility, skills, collaboration capacity, and environmental costs [59,60]. In this context, che*C*k*OVER* embodies a simple yet powerful principle: that complex, multi-layered insights can emerge from the most minimal but accurate datapoint in biodiversity science—the validated geographic occurrence. From these basic, traceable units, the framework constructs interoperable outputs that expand into biogeographic, ecological, and conservation insights of high complexity. This minimalist-to-complex design shows how what might appear as a trivial piece of information—the location of a species at a given point in space and time—can be transformed into a rich knowledge package while retaining fidelity to the original observation.

This approach contrasts with much of traditional biodiversity informatics, where raw occurrence data, spatial products, and analytical results exist in separate silos, often with limited connectivity [61]. che*C*k*OVER* explicitly bridges these divides: species-level narratives, georeferenced distribution layers, and conservation metrics are all produced as linked outputs, maintaining lineage back to the primary evidence. Building on earlier efforts to enrich biodiversity data and harmonize infrastructures [40,62], che*C*k*OVER* advances the field by making outputs not only interoperable but also directly consumable by large language models. Unlike prior initiatives that emphasized standards or database interoperability alone, che*C*k*OVER* packages biodiversity information in formats designed to serve simultaneously human researchers and AI-driven reasoning systems.

### Addressing the provenance crisis and pathways of AI uptake

A defining challenge of AI-mediated science is the erosion of source attribution. LLMs often synthesize information from diverse sources without maintaining clear provenance chains, leading to responses that may be factually accurate but lack verifiable attribution [63–65]. che*C*k*OVER* directly targets this challenge by embedding standardized citation metadata in every output, so that AI systems and other downstream tools can, when appropriately configured, provide not only accurate information but also direct pathways to primary sources. This approach has broader implications for scientific integrity in the AI era. As LLMs become primary interfaces for accessing scientific knowledge [66,67], the ability to trace information back to peer-reviewed publications becomes essential for maintaining trust in science-based decision-making [68,69]. che*C*k*OVER*’s structured outputs enable RAG approaches where AI systems can dynamically access and cite specific biodiversity knowledge packages rather than relying solely on pre-trained information that may become outdated or lack source attribution.

Equally important, however, is understanding how and when biodiversity resources become accessible to AI. Current LLMs are not continuously updated from the web; instead, they integrate new data only through periodic retraining by providers [70]. This means that che*C*k*OVER* outputs will enter the permanent core of LLM knowledge only when included in future training snapshots. In the interim, near real-time interoperability can still be achieved through web indexation and RAG-enabled systems, as well as through direct API endpoints that allow models with web connectivity to retrieve updated outputs on demand. By structuring outputs as JSON packages with embedded provenance, che*C*k*OVER* maximizes its visibility for both long-term model training and immediate dynamic retrieval.

#### Version-aware and incremental updates

Biodiversity occurrence datasets are inherently dynamic, with new records, corrections, and extinction annotations accumulating continuously over time. Recomputing full species-level analyses from scratch at each update becomes increasingly impractical for data-rich taxa, particularly when computationally expensive geospatial operations (e.g., basin overlays, AOO/EOO estimation, map generation) are involved. To address this, che*C*k*OVER* is designed around a version-aware and incrementally updatable architecture.

At the core of this design is the separation between raw occurrence records and derived spatial knowledge units. While point-level occurrences constitute the primary input, che*C*k*OVER* stores and exposes species-level outputs in terms of aggregated, semantically meaningful spatial entities (e.g., hydrographic basins, countries, protected areas), together with associated occurrence counts and temporal metadata. These aggregated units, rather than individual points, represent the stable state of a species known distribution at a given processing version. Each run produces a versioned snapshot that records: (i) the data snapshot date, (ii) the che*C*k*OVER* software version, and (iii) the complete set of spatial units currently occupied by the species, separately for indigenous and non-indigenous records. For each unit, the snapshot retains occurrence counts and, where applicable, extinction annotations. This enables downstream comparisons between successive runs without requiring access to the full historical point dataset.

When new data becomes available, che*C*k*OVER* can operate in an incremental update mode. Instead of reprocessing the entire occurrence dataset, the framework identifies records added or modified since the previous snapshot and performs geospatial enrichment only on this delta subset. Newly intersected spatial units are merged with those stored in the previous snapshot, and occurrence counts are updated accordingly. Conversely, when extinction annotations remove the last known occurrence from a spatial unit, that unit is flagged as absent from the current distribution while remaining available for historical context. Geospatial artefacts such as basin-level distribution maps are regenerated from the updated set of occupied spatial units, rather than recomputed from all raw points. This approach dramatically reduces memory requirements and processing time for large datasets, while preserving full reproducibility and traceability. Historical and current ranges are explicitly distinguished, enabling quantitative assessment of range contraction or expansion across versions.

This incremental, version-aware strategy ensures that che*C*k*OVER* outputs remain scalable as biodiversity datasets grow, supports longitudinal analyses of distributional change, and aligns with FAIR principles by making both data provenance and processing history explicit. Importantly, it allows computationally intensive species (e.g., globally invasive crayfish with tens of thousands of records) to be re-evaluated efficiently as new data accumulate, without compromising methodological consistency or interpretability.

### Scalability and generalizability across taxonomic domains

While our proof of concept focuses on crayfish (Astacidea), che*C*k*OVER*’s modular architecture is designed for broad applicability across taxa. The ingestion module accepts heterogeneous occurrence tables by automatically standardizing commonly used coordinate and date formats (e.g., decimal degrees, DMS, split-degree fields, multiple date conventions) and by recognizing alternative column labels for latitude, longitude, and year. Once harmonized, all datasets are processed through the same internal schema, ensuring that standardized outputs remain consistent across species.

The framework is robust to complex distributional patterns—including multi-continental ranges, invasive populations, and temporally stratified records—making it suitable for marine taxa, terrestrial organisms, fossil occurrences, or trait-based datasets with spatial components. Minor adjustments may be required—for example, alternative vernacular-name sources or modified distributional thresholds—but the workflow and output architecture remain unchanged. This design philosophy balances technical flexibility with standardized outputs, enabling interoperability across biological domains while preserving taxon-specific nuance [71,72].

### Scientific impact and institutional applications

The framework’s relevance for conservation and policy is immediate. IUCN Red List assessments, for example, rely on metrics to evaluate extinction risk [50]. che*C*k*OVER* automates these calculations with explicit bibliographic grounding, offering more efficient and consistent workflows across taxonomic groups. In addition to standard range metrics, the framework also reports simple distributional fragmentation signals for endemic and regional taxa, offering an early indication of whether known localities form cohesive or spatially discontinuous units. While intentionally minimalistic, these signals provide useful first-pass context for connectivity considerations and flag cases where more detailed ecological or genetic assessments may be warranted. The framework’s explicit separation of indigenous and non-indigenous distributions supports multiple analytical workflows. For species assessment under IUCN criteria, only native range metrics are biologically meaningful—che*C*k*OVER* ensures these are computed exclusively from indigenous records. For invasion biology applications, comprehensive tracking of alien populations enables researchers to identify biosecurity priorities and evaluate management interventions using the separately computed non-indigenous metrics. This dual-output design allows the same occurrence dataset to serve conservation assessment and invasion monitoring workflows without methodological compromise. Similarly, protected area agencies could leverage dynamically updated species packages to identify coverage gaps, monitor range shifts, or prioritize management interventions based on up-to-date knowledge. This capacity is particularly critical for freshwater taxa, where limited benefits were noted due to their specific requirements on conservation goals [73].

Beyond institutional contexts, che*C*k*OVER* is designed to open biodiversity knowledge to wider audiences. Once its outputs are exposed through user-facing interfaces or AI-mediated tools, non-specialists (including educators, journalists, and citizens) could obtain scientifically grounded answers to basic questions about species. Crucially, each output links back to primary sources, providing any downstream system with explicit citation pathways and countering the drift toward derivative summaries that accumulate errors across successive retellings. In this way, che*C*k*OVER* has the potential to foster trust and engagement with biodiversity science outside academic circles.

### Future developments and broader applications

The modular architecture of che*C*k*OVER* enables several natural extensions that could significantly expand its analytical scope. Integration with trait databases, molecular repositories, and ecological interaction networks would support multidimensional species profiles that combine distributional knowledge with functional, phylogenetic, and ecological context. Multi-species analyses represent another promising direction, generating narratives that capture community-level patterns such as co-distributions, predator-prey dynamics, or pathogen-host associations within the same standardized framework.

User-customizable applications offer immediate practical value for research and conservation communities. Researchers could upload occurrence datasets to benchmark sampling effectiveness, identify geographic coverage gaps, or evaluate survey completeness against known distributions. Conservation organizations could integrate monitoring data with che*C*k*OVER* baselines to assess management outcomes, track population trends, or prioritize intervention strategies based on updated distributional knowledge.

A particularly compelling direction builds on the integrated distributional status summary, which consolidates standardized geographic indicators such as EOO, AOO, and the number of known and extinct-population localities, while also incorporating temporal signals when available. The per-species geospatial layers exported by che*C*k*OVER* are explicitly designed as reusable inputs for downstream analytical tools capable of deriving more advanced fragmentation or range-disjunction indicators. The preliminary fragmentation signals produced for endemic and regional taxa further create a natural bridge toward such advanced workflows [74], offering early pointers to potential spatial breaks that downstream tools may refine.

For taxa of conservation concern, such implementations could incorporate custom, time-restricted analyses which—together with other dedicated outputs such as ecological, molecular, or threat overlays—can generate tailored range summaries that more precisely contextualize distributional patterns. These refinements would strengthen the utility of che*C*k*OVER* outputs in conservation workflows by providing clearer evidence of spatial patterns relevant to Red List criteria [50]. While not a substitute for formal, expert-driven IUCN assessments, such standardized and provenance-rich summaries can support early-stage prioritization, help flag species exhibiting spatial decline or emerging fragmentation, and supply consistent, interoperable inputs that streamline subsequent evaluation efforts [75,76].

## Conclusion

che*C*k*OVER* establishes a new paradigm for biodiversity informatics: transforming minimal yet validated datapoints—species occurrence records—into comprehensive, AI-interoperable knowledge products that serve both human experts and computational systems. By systematically converting occurrence data into structured geo-narratives, quantitative metrics, and machine-readable outputs, the framework repositions biodiversity databases from passive archives into active knowledge engines capable of interfacing with emerging AI technologies.

Three technical innovations distinguish che*C*k*OVER* from existing biodiversity infrastructures. First, the framework implements strict stratification of indigenous and non-indigenous occurrence records, enabling simultaneous support for IUCN-aligned conservation assessments and invasion biology applications without methodological compromise. Second, all outputs embed standardized citation metadata and provenance chains, directly addressing the attribution crisis in AI-mediated science by ensuring that biodiversity knowledge remains traceable to peer-reviewed sources even when accessed through LLM intermediaries. Third, the version-aware architecture enables incremental updates and longitudinal tracking, allowing outputs to evolve dynamically as datasets expand while maintaining full reproducibility and historical context.

The framework’s modular, taxon-agnostic design ensures broad applicability beyond the crayfish proof-of-concept. Any taxonomic group with occurrence records can be processed through the same standardized pipeline, generating consistent outputs regardless of data density, geographic extent, or distributional complexity. By integrating multilingual vernacular names and delivering outputs in multiple interoperable formats (JSON, Markdown, KML/KMZ, GeoJSON), che*C*k*OVER* expands access beyond traditional research communities to encompass educators, policymakers, conservation practitioners, and local communities whose ecological knowledge is encoded in vernacular terminology.

As biodiversity faces accelerating threats from global change, the need for rapid, accurate, and traceable knowledge synthesis has never been more urgent. che*C*k*OVER* demonstrates that occurrence repositories—often dismissed as simple data archives—can function as sophisticated knowledge infrastructure when coupled with appropriate computational frameworks. Rather than treating AI as external to biodiversity science, the framework illustrates how specialized scientific communities can proactively shape the integration of their knowledge domains into AI-mediated systems, ensuring that biodiversity information remains authoritative, verifiable, and accessible across diverse user communities and technological platforms.

The open-source implementation, containerized deployment architecture, and comprehensive documentation position che*C*k*OVER* for adoption by biodiversity platforms, research institutions, and conservation organizations worldwide. Because all outputs are fully data-driven, they represent reproducible snapshots of available occurrence data rather than static assessments, updating naturally as datasets grow and scientific understanding evolves. This living-systems approach aligns with the dynamic nature of biodiversity knowledge itself, ensuring that che*C*k*OVER* outputs remain relevant and actionable as global monitoring efforts intensify and new occurrence records accumulate.

## Supporting information

archived supplement files

## Availability of source code and requirements

Project name: che*C*k*OVER*

Project homepage: https://world.crayfish.ro/

Source code repository: https://github.com/dadvilfed/checkcover/, together with a demonstration dataset (RollData) containing sample crayfish occurrence data, canonical species geo-narrative template (File S1), and the offline lookup tables required in the workflow (provided in TSV format to ensure full Unicode compatibility and to support machine-readable reuse): vernacular names expert supplement (Table S2), hydronyms (Table S3), and freshwater ecoregions (Table S4)

Operating system(s): cross-platform (Windows 10/11, Linux Ubuntu 20.04+, macOS 11+)

Programming language: R (≥4.5.2)

Other requirements: pandoc (≥2.0), GDAL (≥3.0), PROJ (≥7.0) libraries for spatial operations

License: GPL-3

Any restrictions to use by non-academics: none

RRID: [not yet assigned, all persistent identifiers will be registered upon final release]

bio.tools ID: [not yet assigned]

Local installation requirements: For typical use cases (datasets with <10,000 records per species): 16 GB RAM, 4 CPU cores, 100 GB disk space. For large-scale applications (>20,000 records per species or processing hundreds of taxa): 256+ GB RAM, 16+ CPU cores, 500+ GB disk space recommended.

Server deployment: RESTful API endpoints for species packages and search functionality, MariaDB for metadata indexing, CDN integration for global accessibility, Docker containerization supported

## Additional files

**Supplementary Table S1.** World of Crayfish^®^ Contributors, including full names and institutional affiliations, who provided validated occurrence data, metadata, or region-specific expertise that supported data verification, contextual interpretation, and the completeness of species-level information used in this study.

**Supplementary Table S2.** Expert-curated vernacular names lookup table, supplementing ITIS-sourced names and providing multilingual coverage.

**Supplementary Table S3.** Hydronym lookup table, linking HydroBASINS unit IDs (levels 6, 8, and 10) to their assigned hydronyms, derived by spatial intersection of basin polygons with named river-line datasets.

**Supplementary Table S4.** Lookup table for freshwater ecoregions, linking FEOW unit IDs to their standardized ecoregion names.

**Supplementary Table S5.** Software dependencies used by the che*C*k*OVER* pipeline. All packages are automatically installed during the standard workflow execution, except for the auxiliary datasets listed in Supplementary Tables S2–S4.

**Supplementary File S1.** Canonical che*C*k*OVER* species geo-narrative template used for standardized human- and LLM-readable outputs.

## Abbreviations

AI: artificial intelligence
AOO: area of occupancy
API: application programming interface
CSV: comma-separated values
EOO: extent of occurrence
FAIR: Findable, Accessible, Interoperable, Reusable
FEOW: Freshwater Ecoregions of the World
IUCN: International Union for Conservation of Nature
JSON: JavaScript Object Notation
KML/KMZ: Keyhole Markup Language / Keyhole Markup Language Zipped
LLM: large language model
MRBW: Major River Basins of the World
RAG: retrieval-augmented generation
TEOW: Terrestrial Ecoregions of the World
TSV: tab-separated values
WDPA: World Database on Protected Areas
WoC: World of Crayfish^®^ platform

## Consent for publication

This study involves species-occurrence data sourced from community contributors, following the anonymization protocol of the WoC^®^ platform [10], ensuring that no precise coordinates are released in any che*C*k*OVER* output. All authors, including WoC^®^ Contributors, have reviewed and approved the final manuscript and consent to its publication.

## Acknowledgments

We thank Marian Neagul (West University of Timi□oara) for configuring and maintaining the server environment hosting the dynamic data platform, and Alexandru E. Mizeranschi (Research and Development Station for Bovine, Arad, Romania) for technical advice during pipeline testing.

## Author contributions

L.P.: conceptualization, investigation, methodology, supervision, validation, writing—original draft. D.L.: software, methodology, formal analysis, visualization, writing—original draft. V.I.B.: methodology, visualization, writing—original draft. C.I.N.: methodology, visualization, writing—original draft. T.T.□.: methodology, visualization, writing—original draft. A.V.L.: methodology, visualization, writing—original draft. WoC^®^ Contributors: supervision, validation. All: writing—review & editing.

## Use of AI tools

ChatGPT 5.1 model [77] was used for language refinement of the manuscript. No scientific interpretations were generated by AI. All content was reviewed and fully validated by the authors, who take full responsibility for the final text.

## Funding

This work received no project-based or grant funding. The Article Processing Charge was supported through doctoral publication funds provided by the West University of Timi□oara.

## Data availability

The processed dataset used in this study has been deposited in GigaDB as a static, citable snapshot (DOI to be assigned upon GigaDB deposition, files will be made available to editors and reviewers through GigaDB’s private access mechanism during peer review, and released publicly upon publication). All raw occurrence records originate from the WoC^®^ platform (https://world.crayfish.ro), which functions as a dynamic, continuously updated repository. The platform provides interactive access to current and future versions of the dataset, as well as to all che*C*k*OVER*-generated outputs across subsequent releases.

## Competing interests

The authors declare that they have no competing interests.

## Ethical approval

The study was approved by an institutional review board at the West University of Timisoara under the identifier UVT2025-086385/24.11.2025.

## Notes

https://github.com/dadvilfed/checkcover/

### Competing Interest Statement

The authors have declared no competing interest.

### Summary of Updates

The manuscript was formatted to meet the Giga Science journal specifications.

