## Supplementary material for "che*C*k*OVER*: An open framework and AI-ready global crayfish database for next-generation biodiversity knowledge": archived supplement files: File_S1.md

# Canonical cheCkOVER Geo‑Narrative Template v0.9

## Purpose
Standardized, reproducible template for generating human‑readable and AI‑ready (JSON) biodiversity knowledge outputs within the cheCkOVER framework.

## Scope
Applies to all species‑level packages generated by cheCkOVER.

## Version
0.9 (December 2025)

---

## 1. TAXONOMIC IDENTITY

### Purpose
Establish an unambiguous taxonomic reference.

### Required fields
- **Scientific name:** `{Genus species Author, Year}`
- **Higher taxonomy:** `{Order > Superfamily > Family}`

### Optional fields
- **Common names (multiple languages)**  
  Format: `{Language name}: {name_1; name_2; …}`  
  Example: `English: idle crayfish; Romanian: racul bihorean`

  Notes:
  - Language names are rendered in full (human‑readable form).
  - Internal language identification and mapping are handled using ISO 639‑3 codes within the cheCkOVER workflow.
  - Multiple names per language are allowed.
  - Vernacular names are sourced exclusively from official nomenclatural resources (e.g. ITIS) or expert‑curated scientific literature.
  - Vernacular names do not represent automated or ad hoc translations into different languages.
  - When vernacular names are unavailable, users are encouraged to rely on the English common name (if available) or on the scientific Latin name.

### Conditional fields (display only if available)
- **Type locality**
  - Geographic unit: `{Hydrographic basin name, Country}`
  - Protected area: `{WDPA name}` (if applicable)
  - Bibliographic reference: `{Author, Year, DOI/URL}`

---

## 2. INDIGENOUS RANGE OVERVIEW

### Scope
Computed exclusively from **indigenous** records, defined as occurrences with origin `native` or `type_locality`.

### Purpose
Provide conservation‑relevant distributional metrics consistent with IUCN practice.

### 2.1 Distribution Summary

#### Required metrics
- **Distribution category:** `endemic | regional | cosmopolitan`
- **Extent of occurrence (EOO):** `{value} km²`
- **Area of occupancy (AOO):** `{value} km²`

#### Geographic distribution
- **Countries (native):** `{list ordered by record count, descending}`
  - Count: `{n}`

- **Subnational administrative units (native):** `{list ordered by record count, descending}`
  - Count: `{n}`

- **Hydrographic basins (native):** `{deduplicated list of basin names, list ordered by record count, descending}`
  - Count: `{n}`

#### Conservation context
- **Protected area coverage**
  - Native records within protected areas: `{n} ({percent}%)`
  - Protected areas represented: `{list ordered by record count, descending}`

#### Biogeographic context
- **Terrestrial ecoregions (TEOW):** `{list}`
- **Freshwater ecoregions (FEOW):** `{list}`

#### Temporal context
- **Temporal coverage:** `{year_min}–{year_max}`
- **Recent records:** `{n} records post‑2000 ({percent}%)`

### 2.2 Auto‑Generated Contextual Statements

Generated through rule‑based logic:

- **Endemic:**  
  The species is restricted to a limited geographic range.

- **Regional:**  
  The species is native across a broader regional extent.

- **Cosmopolitan:**  
  The species’ native range spans multiple continents.

- **Protected area coverage <25 percent:**  
  Only a minority of native occurrences fall within protected areas.

- **Protected area coverage ≥75 percent:**  
  Most native occurrences fall within protected areas.

- **Short temporal span:**  
  Occurrence data span a limited temporal window.

- **Few recent records:**  
  Most records predate 2000, suggesting outdated distributional knowledge.

## 2.3 INDIGENOUS RANGE DYNAMICS (Historical Context)

### Display condition
Shown only when historical and current ranges differ due to documented local extinctions.

### Metrics
- **Historical EOO:** `{value} km²`
- **Historical AOO:** `{value} km²`
- **Current EOO:** `{value} km²`
- **Current AOO:** `{value} km²`
- **Range change signal:** `stable | contraction | expansion`
- **Magnitude of change:** `{delta_EOO_percent, delta_AOO_percent}`

### Extinction summary
- **Extinct localities:** `{n}`
- **Extinction timeframe:** `{year_first–year_last}`
- **Regional breakdown:** `{Country: n}`

## 2.4 FRAGMENTATION ASSESSMENT (Indigenous Range Only)

### Scope
Applied only to endemic and regional species.  
Cosmopolitan species are excluded by design.

### Method summary
Adaptive spatial clustering of indigenous localities in an equal‑area projection.

### Output
- **Fragmentation signal:** `detected | not_detected`

If detected:
- **Number of clusters:** `{n}`
- **Cluster composition:** `{cluster_id: n_records (percent)}`
- **Interpretation:**  
  Native occurrences are distributed across multiple spatially disjunct clusters.

### Note
This is a conservative descriptive signal and does not represent a formal connectivity or Red List assessment.

---

## 3. NON‑INDIGENOUS RANGE OVERVIEW (Informative)

### Scope
Computed from records with occurrence origin `alien`, `introduced`, or `cryptogenic`.

### Purpose
Document invasion dynamics and biosecurity context.  
Metrics in this section do not contribute to conservation status assessment.

### 3.1 Status
- **Non‑indigenous populations:** `present | not_detected`
- **Occurrence origin(s):** `alien | introduced | cryptogenic`

### 3.2 Informative Metrics
- **Extent of occurrence (EOO):** `{value} km²`
- **Area of occupancy (AOO):** `{value} km²`
- **Countries:** `{list ordered by record count, descending}`
- **Subnational administrative units:** `{list ordered by record count, descending}`
- **Hydrographic basins:** `{deduplicated list of basin names, list ordered by record count, descending}`
- **Protected areas represented:** `{list ordered by record count, descending}`
- **Temporal coverage:** `{year_min}–{year_max}`
- **First record year:** `{year}`

### 3.3 Non‑Indigenous Range Dynamics (if applicable)
- **Range change signal:** `stable | expansion | contraction`
- **Magnitude of change:** `{delta_EOO_percent, delta_AOO_percent}`

---

## 4. DATA QUALITY, TRACEABILITY, AND PROVENANCE

### Purpose
Document data completeness, reliability, and full provenance of the geo-narrative outputs generated by cheCkOVER.

### 4.1 Data summary
- **Total records analyzed:** `{n_total}`
  - **Indigenous records:** `{n_indigenous}`
  - **Non-indigenous records:** `{n_non_indigenous}`
- **High-accuracy records:** `{n_high_accuracy} ({percent}%)`
- **Data quality score:** `{0–100}`  
  *(Composite indicator reflecting spatial accuracy, temporal precision, and source reliability)*

### 4.2 Bibliographic coverage
- **Total references:** `{n}`
- **DOI-linked references:** `{n} ({percent}%)`
- **Bibliography handling:**  
  Full, de-duplicated bibliography of raw data sources provided as a separate artifact and linked to this geo-narrative output.

### 4.3 Raw data provenance
- **Raw data repository:** World of Crayfish®
- **Raw data citation:**
  Ion et al. (2024)
  *World of Crayfish™: A web platform towards real-time global mapping of freshwater crayfish and their pathogens*
  *PeerJ 12. e18229*
  https://doi.org/10.7717/peerj.18229

### 4.4 Processing framework provenance
- **Processing framework:** cheCkOVER
- **Software citation:**
  Pârvulescu et al. (2026)
  *cheCkOVER: An open framework and AI-ready global crayfish database for next-generation biodiversity knowledge*
  *GigaScience*
  https://doi.org/[to be assigned]
- **cheCkOVER version:** `{version}`
- **Processing date:** `{ISO date}`
- **Data snapshot date:** `{YYYY-MM-DD}`

### 4.5 Interpretation note
This geo-narrative is a **descriptive synthesis** derived from curated occurrence data and standardized spatial analyses.  
It does **not** constitute a formal conservation status assessment and should be interpreted in conjunction with primary sources and expert evaluation.

---

## 5. FORMAL NARRATIVE SUMMARY (Human‑Readable)

### Purpose
Produce a continuous, publication‑ready synthesis of approximately 300–500 words, suitable for encyclopedic, policy, and scientific contexts. The narrative should be accessible to both domain experts and general audiences, while maintaining scientific rigor and traceability.

### Design principles
- **Logical flow:** Progress from taxonomic identity → geographic context → conservation status → data quality
- **Contextual integration:** Weave quantitative metrics into narrative prose rather than listing them
- **Clarity and precision:** Use specific numbers and named geographic units; avoid vague statements
- **Transparency:** Distinguish between native and non-indigenous ranges; flag data limitations when present
- **Actionability:** Highlight conservation-relevant signals (fragmentation, protected area coverage, extinction events)

### Content structure and guidance

#### **Paragraph 1: Taxonomic identity and distributional category (50–80 words)**

**Components to include:**
- Scientific name with author citation
- Higher taxonomy (Order, Family)
- Common names (select 1–2 most widely used, with full language names)
- Distributional category (endemic/regional/cosmopolitan) with supporting evidence
- Type locality (if available), linked to bibliographic source

**Narrative guidance:**
Open with the species' taxonomic identity and immediately contextualize its biogeographic status. For endemic species, emphasize restricted range; for cosmopolitan species, note broad distribution. If vernacular names exist, include them with language names written in full (e.g., "English", "română", "Deutsch") to improve accessibility.

**Example opening:**
> *Euastacus bispinosus* Clark, 1941 (Decapoda: Parastacidae), commonly known as the Glenelg spiny crayfish (English), is an **endemic** freshwater species restricted to southeastern Australia. With a native extent of occurrence of 6,965 km², the species' limited geographic footprint places it within restricted-range taxa of conservation concern.

#### **Paragraph 2: Indigenous range—geography and biogeography (100–120 words)**

**Components to include:**
- Countries and key subnational units (top 2–3 by record count)
- Hydrographic basins (emphasize primary basins, mention count)
- Ecoregions (TEOW/FEOW) for ecological context
- AOO value (if significantly smaller than EOO, note the discrepancy)
- Temporal coverage (first and last record years; percentage of recent records)

**Narrative guidance:**
Describe WHERE the species occurs using named geographic and hydrological units. Integrate biogeographic context (ecoregions) to situate the species ecologically. If the species spans multiple countries or dozens of basins, summarize hierarchically (e.g., "recorded across 12 basins within the Danube drainage system"). Flag temporal biases if <25% of records are post-2000.

**Example:**
> Native occurrences span **7 hydrographic basins** including the Glenelg River, 8 Mile Creek, and Drain 88, all within southeastern Australia. The species inhabits the **Naracoorte woodlands** and **Southeast Australia temperate forests** terrestrial ecoregions, and is associated with Freshwater Ecoregion 915. Validated records span **[year_min]–[year_max]**, with **[X]% collected post-2000**, indicating [adequate/limited] recent survey coverage.

#### **Paragraph 3: Conservation context—protected areas, fragmentation, and extinction signals (80–100 words)**

**Components to include:**
- Protected area coverage (percentage and key protected areas by name)
- Fragmentation signal (if detected: number of clusters, relative sizes, interpretation)
- Extinction events (if present: count, timeframe, affected regions)
- Conservation interpretation (integrate signals with IUCN-aligned language where appropriate)

**Narrative guidance:**
Assess the species' conservation standing using DESCRIPTIVE language (avoid formal Red List categorization unless already assessed). If fragmentation is detected, explain its conservation implications (e.g., "native occurrences form three spatially disjunct clusters, raising concerns about population connectivity"). For species with extinction records, quantify the loss and note temporal trends (e.g., "13% of documented extinctions occurred post-2000").

**Example:**
> Only **35.3% of native records** fall within designated protected areas, including Glenelg Estuary and Discovery Bay Wetlands, indicating limited formal conservation coverage. Distributional analysis reveals **fragmentation**: native occurrences form **3 spatially disjunct clusters** (76%, 12%, 12% of localities), suggesting potential barriers to gene flow and heightened vulnerability to localized threats. [If applicable: *X* extinct localities documented between [year]–[year], concentrated in [region].]

#### **Paragraph 4: Non-indigenous populations (if present) (50–70 words)**

**Display condition:** Only include if non-indigenous records exist (alien/introduced status).

**Components to include:**
- Non-indigenous status (alien/introduced/cryptogenic)
- Countries and basins where introduced
- EOO/AOO of alien range (compare to native range if informative)
- First introduction year (if known)
- Temporal trend (stable/expanding/contracting)

**Narrative guidance:**
Clearly mark this as INFORMATIVE context that does not affect native-range conservation metrics. Use language like "Outside its native range..." or "Introduced populations have been documented in...". If the alien range exceeds the native range, flag this explicitly.

**Example:**
> **Non-indigenous populations:** No alien or introduced occurrences documented. [OR] Outside its native range, introduced populations are established in **[countries]**, spanning **[X] hydrographic basins** and representing an alien EOO of **[X] km²**. The earliest introduction record dates to **[year]**, with evidence of **[stable/expanding]** range dynamics since.

#### **Paragraph 5: Data quality, provenance, and interpretation caveats (60–80 words)**

**Components to include:**
- Total records analyzed (native + alien breakdown)
- High-accuracy records (percentage)
- Data quality score (if computed)
- Bibliographic coverage (total references, DOI-linked percentage)
- Provenance statement (raw data source, processing framework version)
- Interpretation caveat (this is a descriptive synthesis, not a formal Red List assessment)

**Narrative guidance:**
Conclude with transparency about data completeness and quality. Acknowledge limitations (e.g., "Most records lack precise temporal metadata") or strengths (e.g., "High-accuracy coordinates available for 89% of occurrences"). Always include the interpretation caveat to prevent misuse as a formal assessment.

**Example:**
> This synthesis is based on **[X] validated occurrence records** ([X]% native, [X]% non-indigenous), of which **[X]% meet high spatial accuracy standards** (coordinate uncertainty <1 km). The analysis incorporates **[X] bibliographic references** ([X]% DOI-linked), processed through cheCkOVER v[X.X] on [date]. Raw data are maintained in the World of Crayfish® repository (Ion et al. 2024). **This geo-narrative is a descriptive synthesis and does not constitute a formal IUCN Red List assessment.**

### Output format
- Render as continuous prose (no section headers within the narrative)
- Integrate citations inline (e.g., "Ion et al. 2024" or "cheCkOVER v1.0")
- Use **bold** for key conservation signals (endemic status, fragmentation detected, extinction events)
- Maintain consistent past tense for historical data, present tense for current status

### Quality checklist
Before finalizing, verify:
- [ ] All numeric values are non-placeholder and match computed metrics
- [ ] Geographic units are named (not just counted)
- [ ] Names used for all geographic/administrative units (not codes or IDs)
- [ ] Temporal context is explicit (year ranges, percentage recent records)
- [ ] Conservation signals are integrated, not merely listed
- [ ] Non-indigenous section is clearly demarcated (if present)
- [ ] Provenance and interpretation caveat are included
- [ ] Narrative reads fluidly without bullet points or excessive formatting

---
